# Signature morpho-electric, transcriptomic, and dendritic properties of extratelencephalic-projecting human layer 5 neocortical pyramidal neurons

**DOI:** 10.1101/2020.11.02.365080

**Authors:** Brian E. Kalmbach, Rebecca D. Hodge, Nikolas L. Jorstad, Scott Owen, Trygve E. Bakken, Rebecca de Frates, Anna Marie Yanny, Rachel Dalley, Lucas T. Graybuck, Tanya L. Daigle, Cristina Radaelli, Matt Mallory, Medea McGraw, Nick Dee, Philip R. Nicovich, C. Dirk Keene, Ryder P. Gwinn, Daniel L Silbergeld, Charles Cobbs, Jeffrey G Ojemann, Andrew L Ko, Anoop P Patel, Richard G. Ellenbogen, Staci A. Sorensen, Kimberly Smith, Hongkui Zeng, Bosiljka Tasic, Christof Koch, Ed S. Lein, Jonathan T. Ting

**Affiliations:** Allen Institute for Brain Science, Seattle, WA; Department of Physiology and Biophysics, University of Washington, Seattle, WA; Dept. of Pathology, Univ. of Wash., Seattle WA USA; Epilepsy Surgery and Functional Neurosurgery, Swedish Neuroscience Institute, Seattle, WA; Dept. of Neurological Surgery and Alvord Brain Tumor Center, Univ. of Wash., Seattle WA USA; The Ben and Catherine Ivy Center for Advanced Brain Tumor Treatment, Swedish Neuroscience Institute, Seattle, WA; Department of Neurological Surgery, University of Washington School of Medicine, Seattle, WA; Regional Epilepsy Ctr., Harborview Med. Ctr., Seattle WA USA

## Abstract

In the neocortex, subcerebral axonal projections originate largely from layer 5 (L5) extratelencephalic-projecting (ET) neurons. The highly distinctive morpho-electric properties of these neurons have mainly been described in rodents, where ET neurons can be labeled by retrograde tracers or transgenic lines. Similar labeling strategies are not possible in the human neocortex, rendering the translational relevance of findings in rodents unclear. We leveraged the recent discovery of a transcriptomically-defined L5 ET neuron type to study the properties of human L5 ET neurons in neocortical brain slices derived from neurosurgeries. Patch-seq recordings, where transcriptome, physiology and morphology are assayed from the same cell, revealed many conserved morpho-electric properties of human and rodent L5 ET neurons. Divergent properties were also apparent but were often smaller than differences between cell types within these two species. These data suggest a conserved function of L5 ET neurons in the neocortical hierarchy, but also highlight marked phenotypic divergence possibly related to functional specialization of human neocortex.

## Introduction

Understanding how cellular diversity relates to cell types and circuits remains one of the biggest challenges in modern neuroscience (Zeng and Sanes, 2017). Within the neocortex, excitatory pyramidal neurons display an astonishing diversity in gene expression, morphology, physiology and response to neuromodulation (Dembrow and Johnston, 2014; Gouwens et al., 2019, 2020; Markram et al., 2015; Sugino et al., 2006; Tasic et al., 2018). One seminal discovery shed light on the organizational structure of these excitatory neuron populations by demonstrating that they can be segregated based on their long-range axonal targets (Harris and Shepherd, 2015). This is exemplified in layer 5 where pyramidal neurons can be broadly segregated into two classes based upon whether their long-range axons project only within the telencephalon or both within and outside of the telencephalon (Baker et al., 2018).

Both intratelencephalic-projecting (IT) and extratelencephalic-projecting (ET) neurons send axonal projections within the telencephalon (i.e. the cerebral cortex, basal ganglia, etc.), but only ET neurons send long-range axons to subcerebral targets such as the spinal cord, pons and thalamus. In this way, cortical activity that directly affects subcerebral processing/behavior is routed largely through ET neurons. In rodents, L5 ET neurons possess distinctive morpho-electric properties, gene expression, local synaptic connectivity, long-range afferents and response to neuromodulators (Anastasiades et al., 2018; Avesar and Gulledge, 2012; Brown and Hestrin, 2009; Dembrow et al., 2010, 2015; Groh et al., 2010; Guan et al., 2015; Hattox and Nelson, 2007; Kalmbach et al., 2013; Kawaguchi, 2017; Kim et al., 2015; Mao et al., 2011; Sheets et al., 2011; Sorensen et al., 2015; Tasic et al., 2018). These properties distinguish ET neurons from nearly every other neocortical cell type (Gouwens et al., 2019). Furthermore, emerging evidence indicates that ET neurons display unique firing properties *in vivo* and contribute to different aspects of perception and behavior (Economo et al., 2018; Kim et al., 2015; Li et al., 2015; Rojas-Piloni et al., 2017; Saiki et al., 2017).

Translating these findings, which have been described primarily in rodents, to the human neocortex is complicated by several factors. First, there are major cross-species differences in the gross-laminar organization of the neocortex, the size of neurons, the laminar expression of genes, and the intrinsic membrane properties of homologous cell types (Beaulieu-Laroche et al., 2018; Berg et al., 2020; Hodge et al., 2019; Kalmbach et al., 2018; Mohan et al., 2015; Zeng et al., 2012). Additionally, there are experimental limitations to studying long-range projections in humans and other primates. In rodents, retrograde tracers or viruses can label ET neurons to target them for patch-clamp physiology or other functional studies (Dembrow et al., 2010; Hattox and Nelson, 2007; Tervo et al., 2016), but these approaches are not possible in humans. In addition, ET neurons appear to be relatively more rare in human compared with rodent neocortex (Hodge et al., 2019, 2020).

Recent advances from single cell transcriptomics offer a unique anchor to identify ET neurons in the human neocortex and to study their cellular properties. Here, we leverage these recent advances to address which distinguishing cellular properties of rodent ET neurons are conserved or divergent in the human neocortex.

## Results

### Cross species differences in the relative abundance of putative L5 ET neurons

In mice, single-cell transcriptomics readily distinguishes ET versus IT neurons. Combined retrograde labeling and scRNA-seq (i.e. Retro-seq) has validated this target/transcriptome relationship (Economo et al., 2018; Tasic et al., 2018). Alignment of human and mouse neocortical taxonomies based on RNA-sequencing has enabled strong inference of homologous transcriptomic cell types in the human cortex that correspond to L5 ET neuron types in the mouse (Hodge et al., 2019, 2020). This alignment reveals a striking cross-species difference in the relative abundance of transcriptomically-defined ET neurons. Putative ET neurons comprise only 2-6% of excitatory neurons in L5 of the human neocortex compared to 20-30% in mice, depending upon cortical area. It is unclear, however, whether the relative rarity of ET neurons is a unique property of the human neocortex. To address this question, we estimated the relative abundance of ET neurons across mouse, macaque and human temporal cortex by using multiplex fluorescence *in situ* hybridization (FISH) against a conserved L5 ET marker gene (*FAM84B*) and a glutamatergic neuron marker (*SLC17a7*; Figure 1A). In all three species, neurons that expressed both *FAM84B and SLC17a7* were characterized by large pyramidal shaped somata (Figure 1B), a hallmark of L5 ET neurons in rodents (Baker et al., 2018; Oswald et al., 2013). Putative ET neurons were most abundant in mice, followed by macaque and then human temporal cortex (Figure 1C). Thus, there are differences in the relative abundance of L5 ET neurons in the neocortical column across species, likely related to cortical expansion and dramatic increase in IT neuron abundance and cell type diversity.

**Figure 1.**
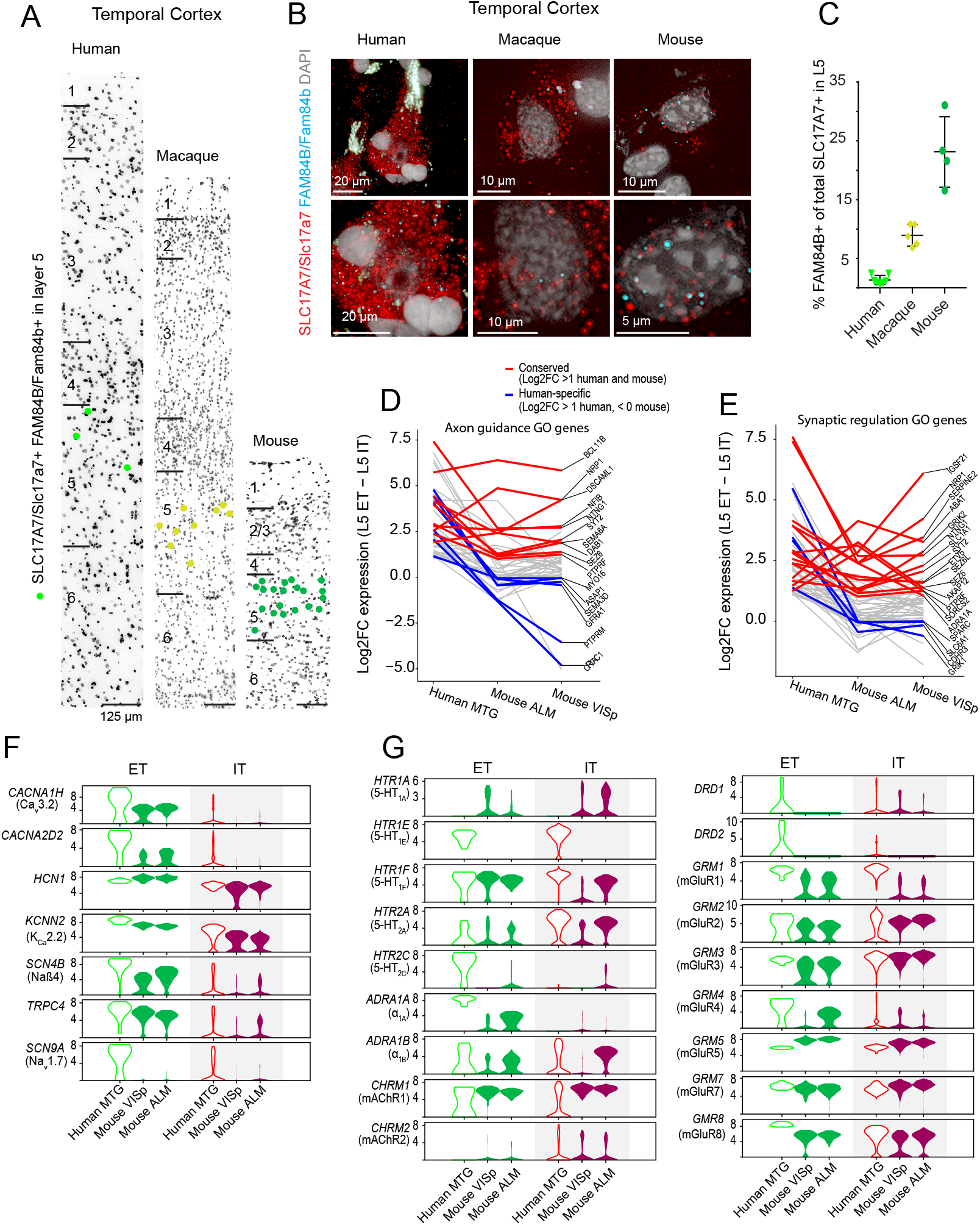
ET neurons are sparse in human MTG, but share many genes with mouse ET neurons A) Representative inverted images of DAPI-stained sections of human, macaque and mouse temporal cortex. Dots denote the location of cells labeled using multiplex fluorescent in situ hybridization (mFISH) for FAM84B and SLC17A7. Horizontal bars denote putative layer boundaries. B) Example mFISH images of FAM84B and SLC17A7 labeling in L5 pyramidal neurons in human, macaque and mouse temporal cortex. C) Quantification of the proportion of SLC17A7 + cells expressing the ET marker FAM84B in temporal cortex of mouse, macaque and human expressed as a fraction of the total number of excitatory cells in L5. Individual data points are denoted by symbols. Line graphs of D) axon guidance and E) synaptic regulation related genes with expression enrichment in L5 ET versus IT neurons in human MTG (> 1 log2 fold-difference) and their respective enrichment in L5 ET neurons in mouse VISp. Notable conserved (red) and human specific (blue) genes are highlighted F) Human L5 ET neuron enriched ion channel genes and their expression in the mouse VISp and ALM. G) Neuromodulator receptor gene expression in human MTG and mouse VISp/ALM.

The identification of a conserved class of transcriptomically defined ET neurons permitted us to identify genes that may contribute to conserved properties of this cell type as well as genes potentially contributing to phenotypic divergence. To identify genes enriched specifically in human ET neurons, we analyzed an existing single nucleus RNA sequencing dataset from human MTG (Hodge et al., 2019). Comparisons across species are complicated by the lack of a clear rodent homologue to human MTG and by evidence that ET neuron gene expression varies greatly between cortical areas in the human and mouse (Hodge et al., 2019; Tasic et al., 2018). Rather than focusing on a single mouse brain region, we therefore utilized a published single cell RNA sequencing dataset derived from two brain regions, the mouse primary visual cortex (VISp) and the anterior lateral motor cortex (ALM). We identified 4,143 genes with at least 0.5 log_2_ fold enriched expression in transcriptomically-defined ET neurons relative to L5 IT neurons in human MTG. Additionally, 477 DE genes were enriched in mouse ET neurons in both VISp and ALM. This gene set was highly enriched for genes associated with axon guidance and synaptic function (Figure 1D,E; red). A notable example includes *BCL11B*, which is required for subcerebral axonal targeting (Arlotta et al., 2005; Canovas et al., 2015). There were also noteworthy examples of human specific ET genes (Figure 1D,E; blue), including *GRIK1*, which encodes the ionotropic glutamate receptor, GluR5. Nonetheless, these data suggest that known phenotypes of rodent ET neurons, such as subcerebral axonal targeting are broadly conserved in the human L5 ET transcriptomic cell type.

In rodents, L5 ET neurons express a unique repertoire of ion channels, which contributes to their specialized physiological properties (Baker et al., 2018). Many of these voltage-gated channels are also targets of neuromodulation. Indeed, the response to various neuromodulators differs between L5 ET and IT neurons in rodents (Dembrow and Johnston, 2014). We therefore identified ion channel and neuromodulator receptor-related genes enriched in transcriptomically-defined L5 ET neurons in the human MTG (Figure 1F-G). Many, but not all, of these genes were enriched in mouse L5 ET neurons in both VISp and ALM. In addition to a major pore forming HCN-channel subunit (*HCN1*), several classes of G-protein-coupled receptors (GPCRs) were also enriched in transcriptomically defined ET neurons in human MTG. Many of these GPCRs are associated with neuromodulators that differentially affect L5 ET versus L5 IT neurons in the rodent. For example, we observed cross-species differences in the expression of genes encoding 5-HT1 and 5-HT2 receptor family subunits (Figure 1G). In mouse L5 ET neurons, *HTR1A* and *HTR1F* were the dominantly expressed subunits whereas in human L5 ET neurons, *HTR1E* (which is absent in the mouse genome) and *HTR1F* were highly expressed, with little *HTR1A* expression. Similarly, *HTR2C* was abundantly expressed in human, but not mouse L5 ET neurons. These data suggest that human and rodent L5 ET neurons might share similar distinctive intrinsic membrane properties and responses to neuromodulation in comparison to neighboring L5 IT neuron types. In contrast, cross-species differences with respect to other human L5 ET enriched genes highlight areas of potential phenotypic divergence.

### Transcriptomically defined L5 ET neurons possess distinctive intrinsic membrane properties

The transcriptomic classification of cell types has proven remarkably predictive of physiological, morphological, and anatomical properties in both human and mouse neocortex (Bakken et al. 2020; Berg et al., 2020, 2020; Economo et al., 2018; Scala et al., 2020; Tasic et al., 2018). We therefore predicted that in human MTG the electrophysiological properties of transcriptomically defined L5 ET neurons would differ from neighboring IT neurons in a manner consistent with previous observations in rodents (Baker et al., 2018; Dembrow et al., 2010; Gouwens et al., 2019). To test this hypothesis, we performed patch clamp recordings from L5 pyramidal neurons in acute brain slices prepared from neurosurgical resections of human MTG. We utilized current injection protocols designed to detect differences in subthreshold and suprathreshold membrane properties. For a subset of experiments we performed Patch-seq analysis in which the nucleus was extracted after concluding whole cell recording protocols and then processed for snRNA-sequencing. From the resulting RNA sequencing data, the expression levels of thousands of genes were used to assign a transcriptomic cell type identity to the physiologically probed cell by mapping to a reference human MTG transcriptomic cell type taxonomy (Figure 2A, see methods for tree-based mapping details). We grouped Patch-seq sampled neurons and non-Patch-seq sampled neurons into physiologically defined types based on their aggregate neurophysiological signatures. We then asked whether physiologically defined neurons corresponded to genetically defined L5 ET and IT neuron types using Patch-seq mapping.

**Figure 2.**
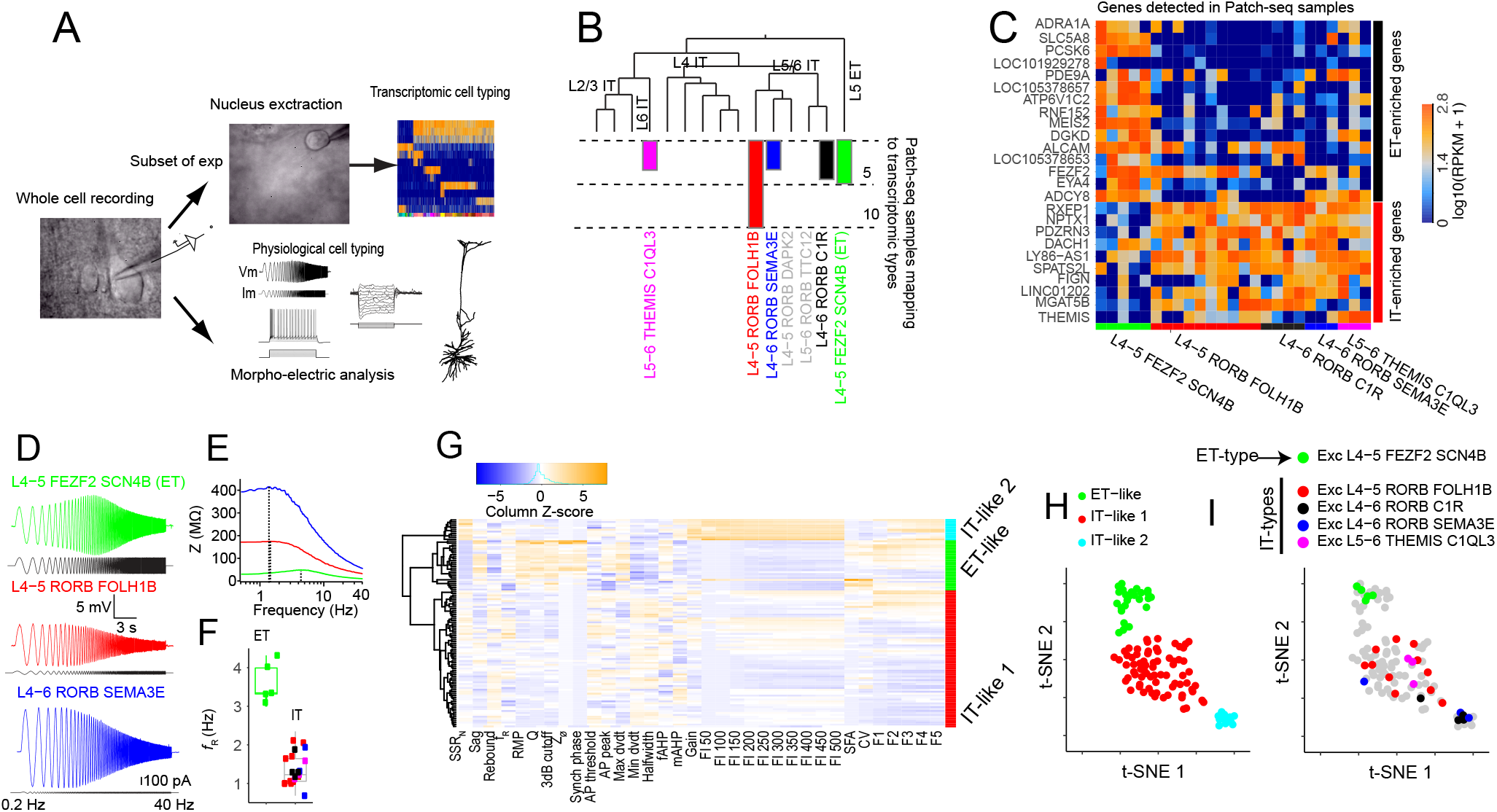
Patch-seq analysis reveals distinctive membrane properties of transcriptomically defined L5 ET neurons. A) We obtained whole cell patch clamp recordings from L5 pyramidal neurons in human brain slices prepared from neurosurgical specimens. For a subset of experiments, Patch-seq analysis was performed in which the RNA content of extracted nuclei were sequenced for the post-hoc assignment of a transcriptomic cell type to the physiologically probed neuron. Several physiological features were extracted from all recordings. B) Glutamatergic transcriptomic cell types in human MTG are shown, with the number of patch-seq samples that mapped with high confidence to each type indicated by the bar plots. C) Heat map of ET and IT enriched genes detected in patch-seq samples. D) Example voltage response of three different transcriptomically-defined cell types to a chirp stimulus and E) Corresponding impedance amplitude profiles (ZAP). Dashed lines denote resonant frequencies. F) Transcriptomically-defined ET neurons displayed a higher resonant frequency than IT neurons (p < 0.001, FDR corrected Mann-Whitney U test). G) Heat map of physiological features of human L5 pyramidal neurons. These features were used to cluster cells into physiologically defined types using Ward’s algorithm. The dendrogram represents the outcome of this clustering. tSNE projection of the features shown in G) color-coded by H) physiologically-defined cell type and I) transcriptomically defined cell type.

Using this approach, we obtained 20 patch-seq samples that mapped with high confidence to one of several infragranular IT types and 5 samples (in green) that mapped to the sole L5 ET cluster in the reference human MTG transcriptomic cell type taxonomy (Figure 2B). As expected, Patch-seq samples mapping to L5 ET and IT types were enriched for multiple established marker genes for the respective types (Figure 2C). Transcriptomically defined L5 ET neurons displayed a higher resonant frequency compared to IT neurons in response to a chirp stimulus (Figure 2D-F), in addition to multiple other pairwise differences in intrinsic properties between transcriptomic cell types (Figure S1). Using unsupervised hierarchical clustering based upon physiology, we observed three distinct types of human L5 neurons (Figure 2G). To visualize the collective differences in the properties of these physiologically defined neuron types we used t-distributed stochastic neighbor embedding (t-SNE) for dimension reduction, where the distance between cells approximates differences in physiological properties and thus cells with similar physiological properties cluster together (Figure 2H). All transcriptomically defined L5 ET neurons belonged to one physiologically defined neuron type whereas all transcriptomically defined L5 IT neurons belonged to one of two different physiologically defined cell types (Figure 2I). For simplicity, we refer to these physiologically defined cell types as ET-like, IT-like 1 and IT-like 2, based on their correspondence to transcriptomic cell types and their broad similarity to rodent L5 ET and IT neurons (see below). Notably, the two IT-like physiologically defined cell types were largely enriched for different L5 IT transcriptomic cell types. Thus, there was a strong correspondence between physiologically and transcriptomically defined cell types.

### Electrophysiological differences in human L5 ET and IT neurons

These results demonstrate that transcriptomically defined L5 ET and IT neurons in human MTG possess distinct electrophysiological properties. To address which physiological features contributed to the clustering of genetically defined L5 ET and IT neuron types, we made pairwise comparisons of specific features between the physiologically defined cell types.

In rodents, subthreshold properties, especially those related to HCN channel expression, readily distinguish L5 ET from IT neurons across several brain regions (Dembrow et al., 2010; Kalmbach et al., 2013; Sheets et al., 2011). Notably, *HCN1*, which encodes a major HCN channel pore forming subunit (Robinson and Siegelbaum, 2003), was enriched in transcriptomically defined L5 ET neurons relative to IT neurons in both mouse and human (Figure 1F). Thus, we predicted that HCN-channel related properties would distinguish L5 ET from IT neurons in human MTG. HCN channels contribute to the resting conductance of a neuron and thus their presence is associated with a lower input resistance and a shorter membrane time constant, resulting in differences in low-pass filtering properties (Kalmbach et al., 2018; Magee, 1998). Additionally, the slow activation and deactivation kinetics of HCN channels contribute to the characteristic voltage sag induced by hyperpolarization and rebound potential upon release from hyperpolarization (Robinson and Siegelbaum, 2003). These kinetic properties endow neurons with membrane resonance in the 2-7 Hz range as well as inductive properties that enable changes in membrane potential to lead changes in current over certain frequencies (Hutcheon et al., 1996; Narayanan and Johnston, 2008; Vaidya and Johnston, 2013). To extract HCN-dependent properties, we measured the voltage response to a series of hyperpolarizing and depolarizing current steps as well as a chirp stimulus (Figure 2A; Figure 3A,B,E).

**Figure 3.**
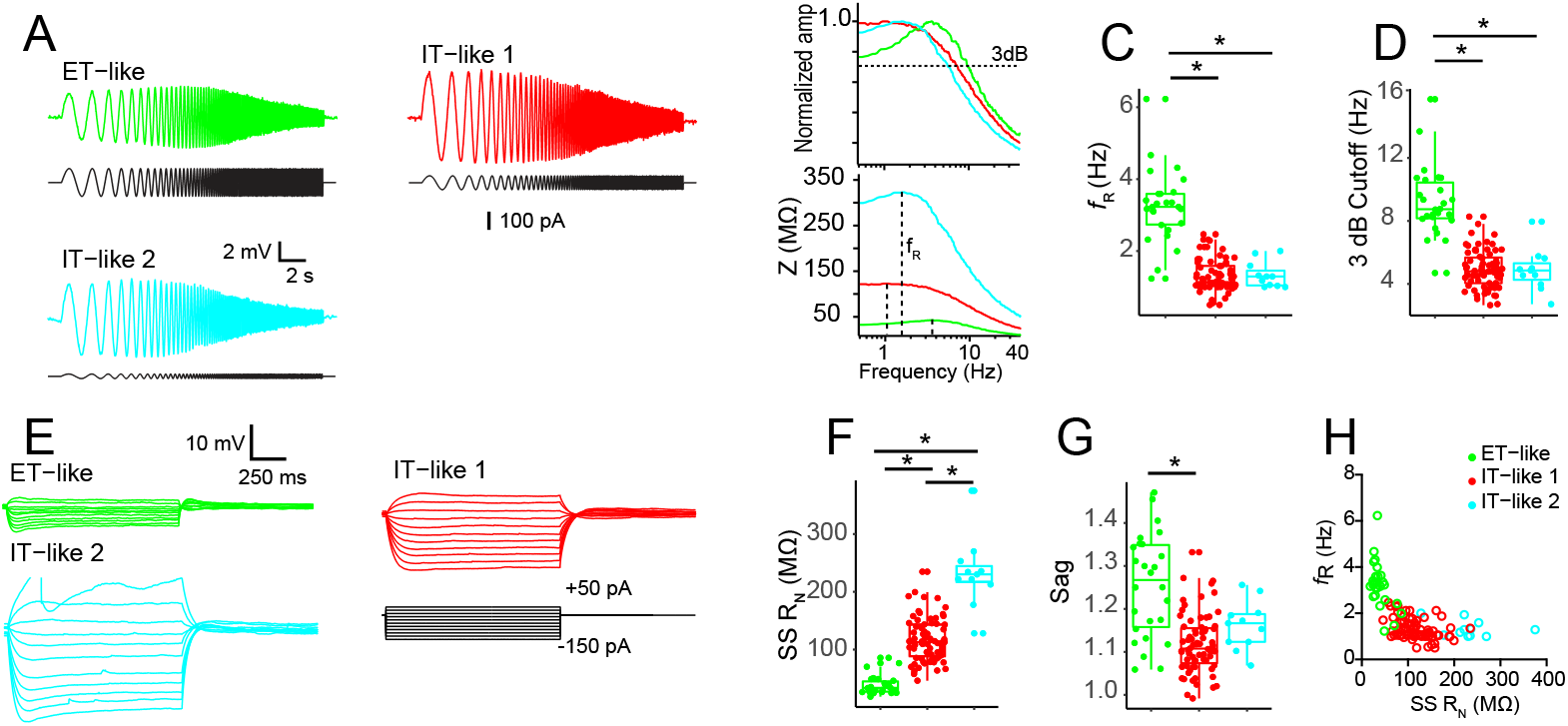
Subthreshold membrane properties of L5 neuron types in human MTG. A) Example voltage response of L5 neuron types to a chirp stimulus. B) ZAP (bottom) and normalized frequency response (top) constructed from the voltage responses in A). Dashed lines indicate resonant frequency (ZAP) and 3dB cutoff (normalized curve). Pairwise comparisons of C) resonant frequency and D) 3 dB cutoff. E) Example voltage responses to a series of hyperpolarizing and depolarizing current injections. Pairwise comparisons of F) Input resistance and G) Sag ratio. H) Resonant frequency as a function of input resistance. * p < .05, FDR corrected Mann-Whitney U test.

We observed several differences in HCN channel-related subthreshold membrane properties between putative L5 ET and IT neurons. ET-like neurons had a higher resonant frequency, higher resonant strength, and higher 3 dB cutoff than both IT-like neuron types (Figure 3C,D; Figure S2). Similarly, the voltage response of L5 ET neurons, but not IT neurons, led the chirp current injection for low frequency components (Figure S2). L5 ET-like neurons also had a lower input resistance and more pronounced voltage sag/rebound than IT-like neurons (Figure 3F-G; Figure S2). Notably, compared with IT-like 1 neurons, IT-like 2 neurons had ~2x higher input resistance and a larger rebound potential. Furthermore, as reported in rodents, simply plotting resonance frequency as a function of input resistance segregated the neurons types well (Figure 3H; Dembrow et al., 2010; Kalmbach et al., 2015). Thus, human L5 ET neurons have more pronounced HCN-related properties than L5 IT neurons.

In addition to subthreshold differences, rodent L5 ET and IT neurons display several known differences in suprathreshold membrane properties (Dembrow et al., 2010; Guan et al., 2015; Hattox and Nelson, 2007; Kalmbach et al., 2013; Oswald et al., 2013; Otsuka and Kawaguchi, 2008; Suter et al., 2013). Action potential properties were extracted from the voltage response to a series of 1 s step current injections of increasing amplitude (Figure 2A). ET-like neurons, perhaps due in part to their low input resistance, displayed the shallowest input/output curves, followed by IT-like and IT-like 2 neuron types (Figure 4A,B). Thus, for a given amplitude current injection, ET-like neurons responded with the lowest number of action potentials. Additionally, we noticed that ET-like neurons tended to respond to near threshold current injections with a high frequency burst of action potentials (Figure 4A), a phenomenon that has been associated with dendritic Ca^2+^ electrogenesis (Beaulieu-Laroche et al., 2018; Larkum et al., 1999; Shai et al., 2015). To illustrate this behavior, we plotted the first instantaneous firing rate (1/first interspike interval) as a function of the amplitude of the current injection above rheobase (Figure 4C). Putative L5 ET neurons displayed the highest instantaneous firing frequencies for at threshold current injections. As an additional way to quantify these bursts, we calculated the percentage of action potentials that occurred within 50 ms of the first spike during the first current step that produced at least 5 spikes. L5 ET-like neurons had the largest percentage of spikes within 50 ms of the first spike and the highest maximum instantaneous firing rate during this current injection (Figure 4D).

**Figure 4.**
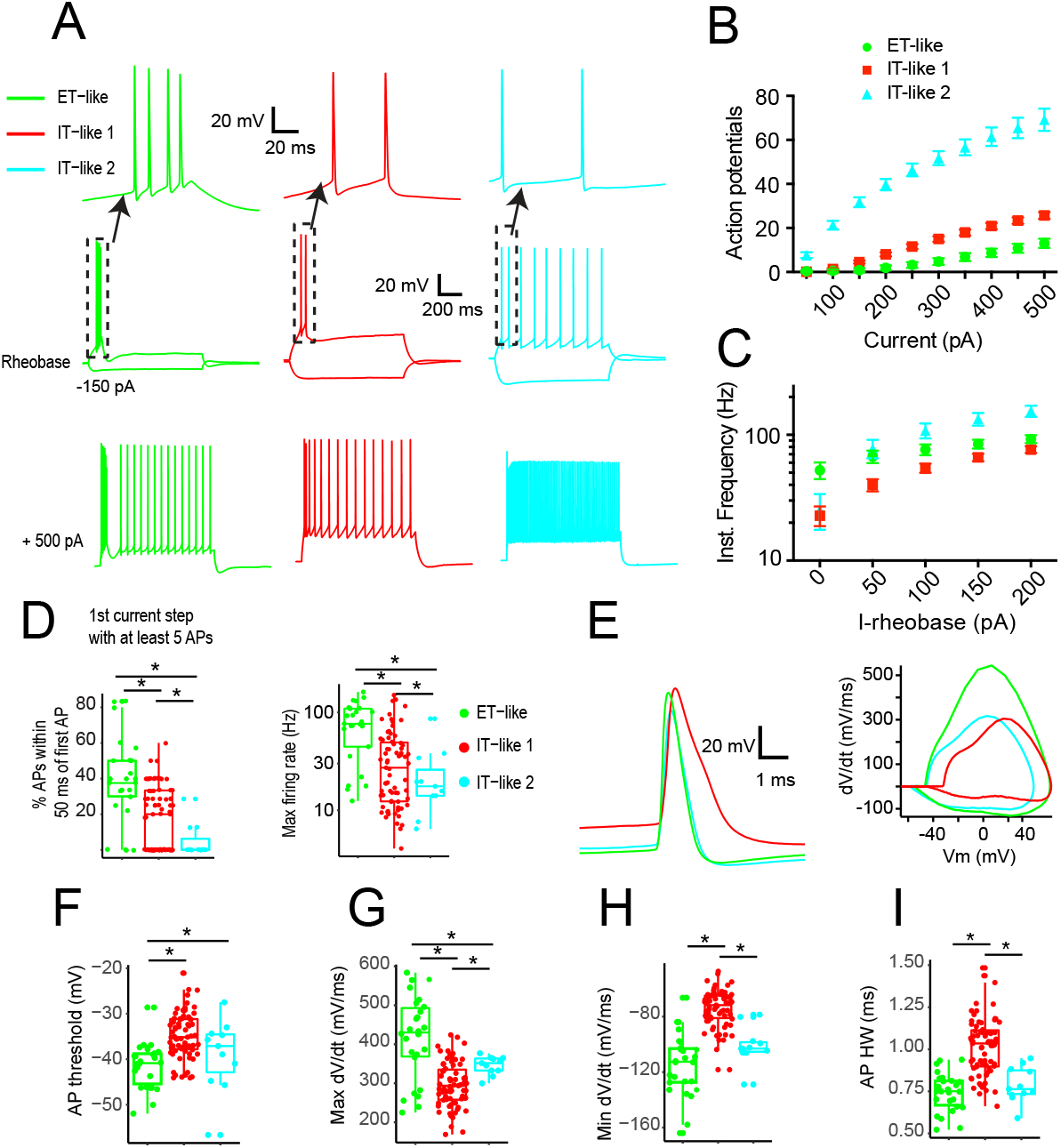
Suprathreshold membrane properties of L5 neuron types in human MTG. A) Example voltage responses to near threshold current injection (middle) and +500 pA (bottom), (top)-expanded view of spikes at near rheobase current injection. B) The number of action potentials evoked as a function of current injection amplitude. C) First instantaneous frequency plotted as a function of current injection amplitude above rheobase. D) (left) The percentage of action potentials occurring within 50 ms of the first spike and (right) maximum instantaneous firing rate for the first current injection producing at least 5 spikes. E) Example action potentials (left) and corresponding phase-plane plots (right). Differences in F) action potential threshold, G) maximum dV/dt H) minimum dV/dt and I) action potential width at half-maximum amplitude. * p < .05, FDR corrected Mann-Whitney U test.

There were several additional differences in action potential properties between putative L5 ET and IT neurons in human MTG that mirror differences observed in rodent neocortex. Example single action potentials and phase-plane plots are presented in Figure 4E. Similar to rodent L5 ET neurons (Dembrow et al., 2010; Pathak et al., 2016; Suter et al., 2013) human L5 ET-like neurons had fast, narrow action potentials characterized by a fast depolarization/repolarization rate, low voltage threshold and narrow width at half-maximum amplitude (Figure 4F-I; Figure S2). Notably, IT-like 2 neurons had similarly fast and narrow action potentials compared to IT-like 1 neurons (Figure 4F-I; Figure S2). ET-like neurons also displayed the largest amplitude medium afterhyperpolarization potentials, and the highest variability in spike timing (Figure S2), perhaps due in part to enriched expression of channels contributing to AHPs and their propensity to burst (Guan et al., 2015). Some of these differences may be explained by differences in the expression of voltage gated ion channels (Bishop et al., 2015; Kalmbach et al., 2015; Miller et al., 2008; Pathak et al., 2016).

As a second approach to investigating which physiological features were most informative to distinguish between L5 ET-like versus IT-like neurons, we trained a series of random forest classifiers on varying subsets of the dataset (Figure S3A). Classifiers had an average accuracy of ~95% when using 40% of the dataset in training and maintained close to this level when using as little as 10% of the dataset. Subthreshold features related to passive membrane properties and HCN conductance had the greatest importance (Figure S3B). The performance of random forest classifiers only marginally decreased when just the top 10 most important features were used in training (Figure S3C). Furthermore, the general shape of t-SNE projections was robust to the removal of several physiological features (Figure S3D,E). Together, these data suggest that subthreshold membrane properties discriminate between L5 ET and IT neurons, but that several other features are sufficient to classify these cell types.

### Putative L5 ET neurons in human MTG have thick-tufted apical dendrites

Classically, rodent L5 ET neurons can be distinguished from L5 IT neurons by their thick apical tuft dendrites (Baker et al., 2018; Dembrow et al., 2010; Gao and Zheng, 2004; Gouwens et al., 2019; Hattox and Nelson, 2007; Oswald et al., 2013). To determine whether human L5 ET-like neurons possess an apical tuft, we performed dendritic reconstructions of physiologically defined neurons. For comparative purposes we also performed dendritic reconstructions of L5 IT-like neurons. Compared with IT-like neurons, ET-like neurons possess a definite apical tuft terminating at the pial surface, but with the tuft branches starting at different distances from the soma (Figure 5A,B). The total length of the apical dendrites (ET 9091.5 ± 559.0 μm, IT 5090.3 ± 759.9 μm; FDR corrected p = 0.026, Mann-Whitney U test) was greater in L5 ET neurons compared with IT-like neurons (Figure 5C). Furthermore, for the apical dendrites, L5 ET-like neurons possessed more dendritic branches (ET 68.7 ± 5.7, IT 34.6 ± 4.8; FDR corrected p = 0.026, Mann-Whitney U test) that on average had a larger diameter (ET 1.04 ± 0.05 μm, IT 0.76 ± 0.06 μm; FDR corrected p = 0.035, Mann-Whitney U test), resulting in greater total surface area (ET 29179.4 ± 1456.8 μm^2^, IT 12601.1 ± 2571.7 μm^2^; FDR corrected p = 0.022, Mann-Whitney U test) compared with IT-like neurons (Figure 5D,E,F).

**Figure 5.**
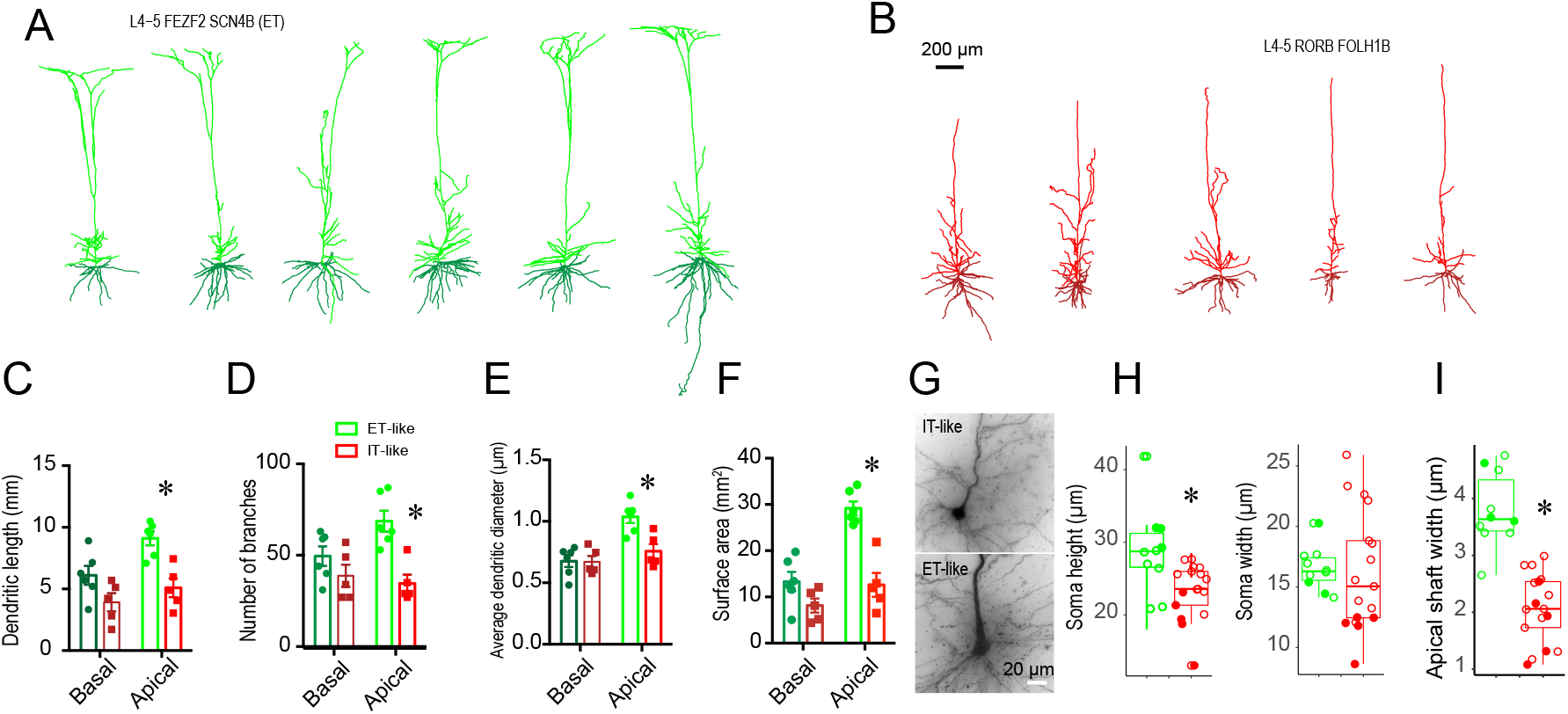
Morphological features of L5 ET- and IT-like neurons in human MTG. Dendritic reconstructions of A) ET-like and B) IT-like neurons in human MTG. Apical and basal dendrites are denoted by different shades of green or red. C) Comparison of basal and apical dendritic length between ET and IT neurons. D) Comparison of the number of basal and apical dendritic branches between ET and IT neurons. E) Comparison of average dendrite diameter between ET and IT neurons. F) Comparison of basal and apical total dendritic surface area between ET and IT neurons. G) Example biocytin fills of perisomatic regions for ET and IT-like neurons. Comparison of H) soma height/width and I) initial apical shaft width. For H and I, filled symbols correspond to transcriptomically-defined cells. * p < 0.05, FDR corrected Mann-Whitney U test.

In addition to dendritic morphology, the somatic morphology of L5 ET and IT neurons is different in rodents. In many cortical areas, L5 ET neuron somata are larger and possess a larger initial apical shaft compared with L5 IT neurons (Gao and Zheng, 2004; Oswald et al., 2013). To quantify somatic shape, we measured the ratio of the somatic height/width as well as the width of the initial apical dendrite shaft of biocytin filled neurons (Figure 5G). In contrast to L5 IT-like neurons, L5 ET-like neuron somata were taller (ET 28.78 ± 1.89 μm, IT 23.31 ± 0.93 μm; FDR corrected p = 0.01, Mann-Whitney U test) than they were wide (ET 16.50 ± 0.56 μm, IT 16.23 ± 1.20 μm; FDR corrected p = 0.47, Mann-Whitney U test; Figure 5H). Similarly, the initial apical shaft was wider in ET-like (3.80 ± 0.21 μm) neurons compared with IT-like neurons (2.07 ± 0.60 μm, FDR corrected p = 0.0001, Mann-Whitney U test; Figure 5I). These properties gave L5 ET-like neuron somata a distinctive teardrop shape that was readily visible in both biocytin fills and under IR-DIC optics during patch-clamp experiments.

### Human L5 ET-like neurons display strong dendritic electrogenesis

To directly test whether the dendrites of human L5 ET neurons display electrogenesis as suggested by their propensity to fire in bursts, we made whole cell recordings from the dendrites of L5 neurons in human MTG (Figure 6A). Example voltage responses to hyperpolarizing and depolarizing current injections measured at different distances from the soma are shown in Figure 6B. For a subset of these dendritic recordings we performed subsequent whole cell recordings from the soma through a second pipette. Projecting the physiological features recorded at the soma onto a tSNE plot revealed that the dendritically probed neurons were L5 ET-like neurons (compare Figure 6C to Figure 2H).

**Figure 6.**
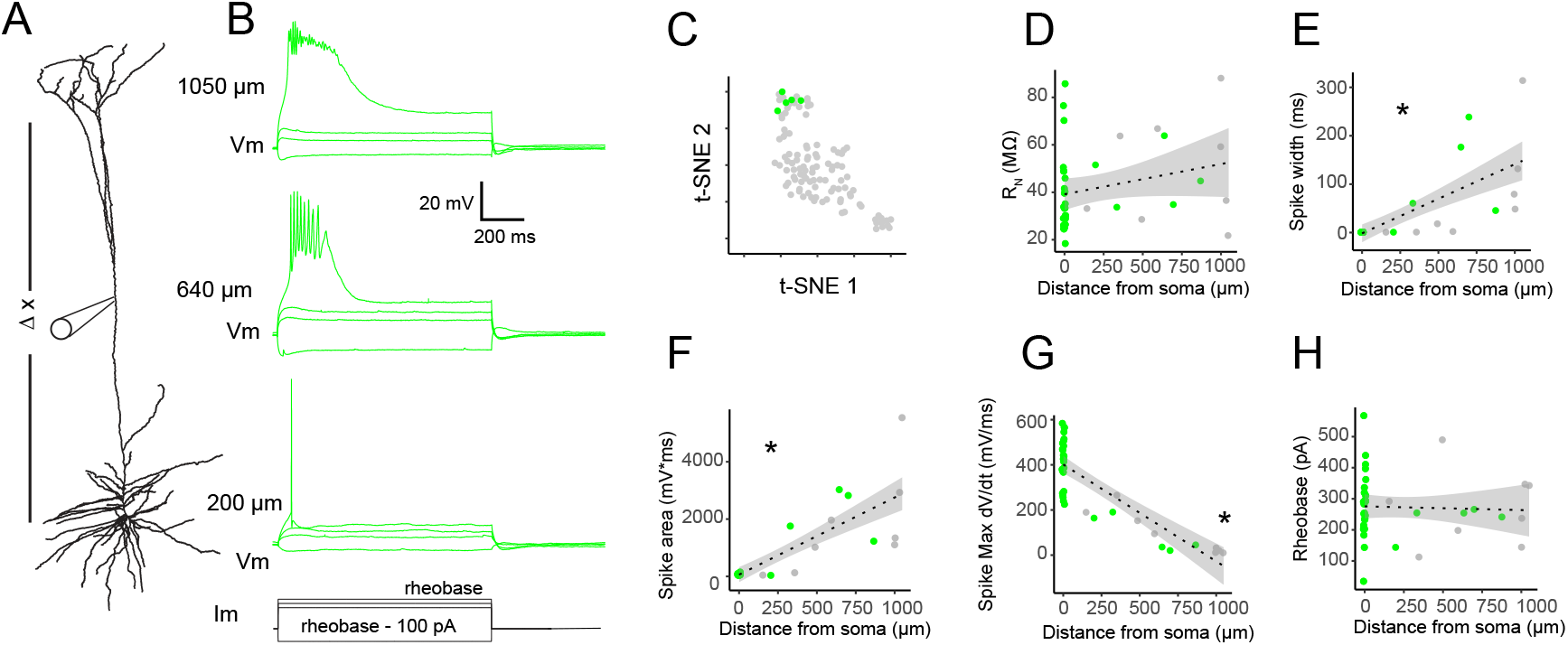
Putative ET neurons display strong dendritic electrogenesis. A) Direct electrical recordings of dendritic membrane properties were performed at various distances from the soma. For a subset of experiments, the soma was subsequently patched with a separate electrode. B) Example voltage responses to hyperpolarizing and depolarizing current injections. Depolarizing current injections were capable of eliciting an all-or-none plateau potential. C) tSNE projection of somatic membrane properties of cells in which the dendritic properties were probed are shaded in green. D) Input resistance (p = 0.13), E) Spike width (p < 0.001), F) plateau potential area (p < 0.001), G) Maximum dV/dt (p < 0.001) and H) Rheobase (p = 0.80) plotted as a function of distance from soma. * p < 0.05, FDR corrected Pearson’s correlation. Shaded region corresponds to SEM. For D-H, green dots denote cells (n = 5) in which the somatic properties were also probed.

We first asked whether the subthreshold membrane properties of human L5 ET-like neurons vary with distance from the soma, as observed in many rodent L5 ET neurons (Dembrow et al., 2015; Kalmbach et al., 2013, 2015, 2017). While input resistance did not change as a function of distance from the soma (Figure 6D), other membrane properties associated with HCN channel expression did. Sag ratio and rebound potential were more prominent at distal recording sites (Figure S4). Furthermore, resonance frequency, 3dB cutoff and total impedance phase increased as a function of recording distance (Figure S4). Thus, as in many rodent L5 ET neurons, HCN related properties increase as a function of distance from soma.

In response to depolarizing current injections, we observed prolonged, all-or-nothing plateau potentials in L5 ET-like dendrites (Figure 6B). The properties of these plateau potentials varied significantly with distance from the soma. The width and area under the curve of the spikes increased with distance from the soma, whereas the maximum rate of rise decreased (Figure 6E-G). Intriguingly, the current required to evoke a plateau potential did not vary with distance from the soma and was similar to the current required to elicit a somatic action potential via somatic current injection. Notably, while dendritic properties were fairly consistent across recordings, we cannot rule out the possibility that some of our recordings were from L5 IT neuron dendrites. Nonetheless, these observations suggest that human L5 ET neurons possess active mechanisms to counteract enhanced compartmentalization associated with their extensive dendritic arbors.

### Cross-cell type variability is greater than cross-species variability

These data suggest that the defining physiological properties of human L5 ET neurons are broadly similar to those reported in rodent neocortex. To directly compare somatic membrane properties across species, we grouped human and mouse L5 neurons into physiologically defined cell types based upon the same physiological features used in Figure 2 (Figure 7A-left). As previously explored with the human data (Figure 2), we then asked whether the physiologically defined cell types corresponded to genetically defined cell types via Patch-seq or labeling by a L5 ET neuron specific enhancer virus (Graybuck et al., 2019). Similar to clustering human data alone, clustering data from both species revealed at least three distinct types of neurons. For both species, physiologically defined L5 ET neurons corresponded to genetically defined L5 ET neurons (Figure 7A-right). Furthermore, ET-like neurons in both species occupied similar areas of the tSNE plot of physiological features (Figure 7A). These data suggest that the intrinsic membrane properties of L5 ET neurons are broadly conserved in mouse and human neocortex.

**Figure 7.**
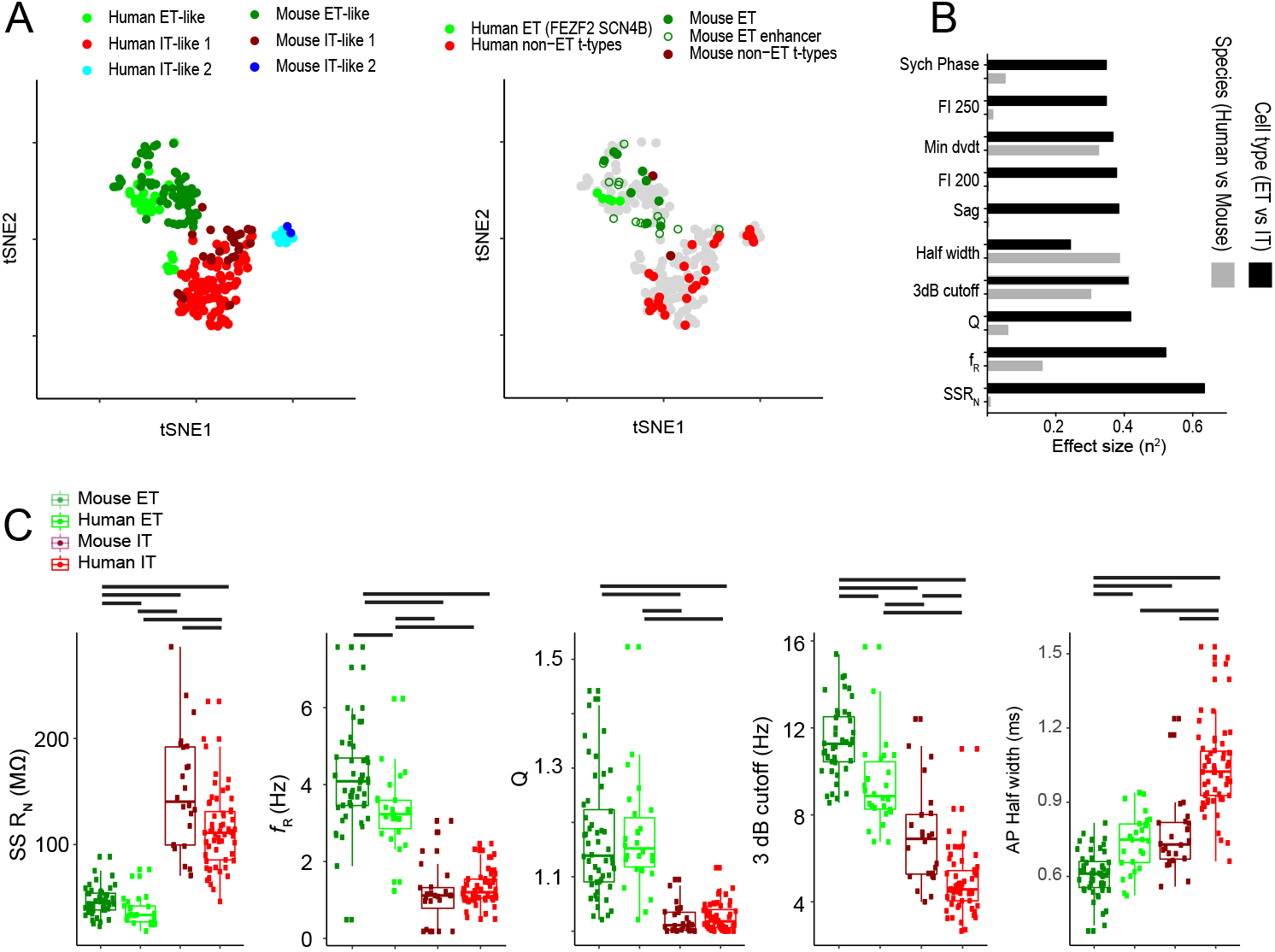
Cross-species comparison reveals conserved and divergent ET neuron somatic membrane properties. A) tSNE of intrinsic membrane properties for human and mouse L5 pyramidal neurons color coded by (left) physiologically-defined cell type and (right) transcriptomic cell type or labeling by an ET-enhancer virus. B) The top 10 largest effect sizes resulting from ANOVA with cell type (ET versus IT) and species (human versus mouse) as the factors. C) Physiological features plotted as a function of species and cell type for the five features with the largest effect sizes. Gray bars denote * p < 0.05, FDR corrected Mann-Whitney U test.

Nonetheless, these observations do not rule out the possibility of subtler cross-species differences in the somatic membrane properties of L5 ET neurons. We thus also made pairwise comparisons between the somatic membrane properties of mouse and human neurons. For this analysis, we collapsed the two IT neuron types into one IT neuron group. We observed both cell type and cross-species differences in nearly every membrane property examined (Figure 7; Figure S5). To determine whether differences are larger between species or between cell types we calculated the effect size associated with each physiological feature. The cell-type-specific effect size was nearly always larger than the effect size determined across species, with a few notable exceptions. For example, cross-species differences were larger than cross-cell-type differences for some suprathreshold properties, including input/output gain and action potential half-width (Figure 7; Figure S5). Taken together, these observations highlight a general conservation of the core defining features of L5 ET and L5 IT neuron types from mouse to human, pointing to a common cell type diversity and resultant functional architecture of L5 across species. These observations also highlight many points of divergence in specific features of L5 ET and IT neuron types between these two species.

## Discussion

In the rodent neocortex, the properties of L5 ET neurons are clearly distinguishable from neighboring L5 IT neurons, which make up the other major class of L5 pyramidal neurons (Baker et al., 2018). Investigation of the distinctive properties of L5 ET neurons in the human neocortex has been hampered by an inability to identify these neurons by their defining feature - long-range subcerebral axonal projections and projection targets. In this study, we circumvent this problem by leveraging transcriptomic definitions of cell types and a mapping of homologous cell types between mouse and human cortical taxonomies. We demonstrate that the morpho-electric properties and gene expression profiles in human L5 ET neurons are as distinctive as their rodent counterparts. Nevertheless, our results reveal substantial divergence and specialization of features.

The identification of a transcriptomic cell type in human middle temporal gyrus (MTG) that is homologous to rodent L5 ET neurons permitted us to identify genes that may contribute to conserved phenotypes (Hodge et al., 2019). In mouse VISp/ALM and human MTG, L5 ET neurons were especially enriched for genes associated with axon guidance and synaptic regulation. These genes may be related to the subcerebral axonal targeting and afferent connectivity of L5 ET neurons. Additionally, both mouse and human L5 ET neurons express different complements of genes encoding neuromodulatory receptors relative to neighboring mouse and human L5 IT neurons. This observation is consistent with the differential effect of neuromodulators on L5 ET and L5 IT neurons in rodent neocortex and suggests human ET neurons may also respond uniquely (with respect to human IT cells) to neuromodulation (Baker et al., 2018; Dembrow and Johnston, 2014). Similarly, rodent and human ET neurons shared assorted differentially expressed ion channel related genes that may contribute to conserved intrinsic membrane properties.

Indeed, patch-seq/patch-clamp recordings revealed broad physiological similarities between rodent and human L5 ET neurons. In both rodent and human L5 ET neurons, HCN-channel-related membrane properties were enhanced compared with L5 IT neurons. In rodents, HCN-channel expression is enriched in the apical dendrite of L5 ET neurons, where it strongly shapes synaptic integration by narrowing the window whereby inputs can be summed in time (Dembrow et al., 2015; Kalmbach et al., 2013, 2015, 2017). These properties tend to make L5 ET neurons most sensitive to inputs containing frequency components in the theta band (4-12 Hz). Our findings indicate that many of these subthreshold integrative properties are conserved in human L5 ET neurons. Human L5 ET neurons also exhibited a distinctive action potential waveform, similar to rodent ET neurons. For example, in both species, action potentials were faster and narrower in L5 ET neurons compared with L5 IT neurons.

Human L5 ET neurons, like rodent L5 ET neurons, tended to respond to near-threshold current injections with bursts of action potentials. In rodent L5 ET neurons, these bursts of action potentials are associated with dendritic Ca^2+^ plateau potentials (Larkum et al., 1999; Shai et al., 2015). Using direct dendritic patch-clamp recordings, we showed that plateau potentials can be elicited in human L5 ET neuron dendrites, and that these events were strongly reminiscent of Ca^2+^ spikes in some rodent L5 neurons. We did not record from confirmed IT neuron dendrites and thus it remains to be seen whether L5 IT dendrites similarly display electrogenesis. Nonetheless, in rodent L5 neurons, plateau potentials serve as a mechanism for associating bottom-up input arriving at the basal dendrites with top-down input arriving at the apical tuft in L1 (Larkum, 2013). Importantly, human L5 ET neurons possessed a pronounced apical tuft in L1, suggesting that they similarly receive substantial feedback input from higher cortical areas. L5 ET neurons in the human neocortex may thus perform a similar computation as rodent L5 ET neurons of coupling feedback and feedforward input in the neocortical circuit.

Our findings confirm some observations from the sole previous report of human L5 dendritic membrane properties (Beaulieu-Laroche et al., 2018), but we also note some key differences. As previously described, HCN-dependent membrane properties varied as a function of distance from the soma, with the exception of R_N_. The flattened somatic-dendritic gradient of R_N_ represents a marked departure from rodent ET neurons (Kalmbach et al., 2013) and may reflect a relative decrease in the density of resting conductance in the dendrite (Beaulieu-Laroche et al., 2018) of human L5 neurons. Unlike (Beaulieu-Laroche et al., 2018), we observed strong electrogenesis in human L5 ET dendrites. While the Beaulieu-Laroche study was performed solely in tissue derived from epileptic patients, we observed dendritic plateau potentials in tissue derived from both epilepsy and tumor patients. Thus, it is unlikely that differences in the underlying medical conditions of patient populations used in these studies contributed to different findings. Aside from methodological differences (pipette solution, external solution, slice preparation, exact surgical sampling etc.), there are several possible explanations for this discrepancy. First, it is unclear exactly which neuronal population(s) were sampled in the previous report. This is especially pertinent given the relative rarity of human L5 ET neurons (2-6% of L5 excitatory neurons in human neocortex versus 20-30% in mouse neocortex (Hodge et al., 2019, 2020; Figure 1). Furthermore, in rodents there is variability in the types of nonlinearities observed in L5 dendrites. For example, the apical trunk of L5 ET neurons in the prefrontal cortex display strong Na^+^ mediated spikes but not Ca^2+^ plateaus, whereas in sensory cortical areas Ca^2+^ electrogenesis is ubiquitous (Gulledge and Stuart, 2003; Harnett et al., 2013; Kalmbach et al., 2017; Larkum et al., 1999; Santello and Nevian, 2015; Shai et al., 2015). Even within a single brain region, there are differences in the propensity of the apical trunk of L5 neurons to generate plateau potentials. Some of this variability appears to be related to differences in dendritic architecture (Fletcher and Williams, 2019). Similar variability in dendritic morphology and related electrogenesis may therefore occur in human neocortex. Nonetheless, we find that at least some human L5 ET dendrites display strong dendritic electrogenesis.

While rodent and human L5 ET neurons had broadly similar genetic and morpho-electric properties, there were nonetheless many cross-species differences that point to potential areas of cross-species divergence and/or specialization. Foremost among these was the striking sparsification of L5 ET neurons from mouse to macaque to human, consistent with previous observations from primary motor cortex (Bakken et al. 2020). This may reflect the dramatic increase in cortical volume relative to subcerebral volume in primates and humans (Heffner and Masterton, 1975; Herculano-Houzel et al., 2015). Additionally, we observed several differences in the expression of genes encoding GPCRs, which yield testable hypotheses concerning cross-species and cross-cell type differences in neuromodulation that could be areas for future study. Genes encoding 5-HT receptors were especially divergent across species, suggesting that human L5 ET neurons may respond differently to serotonin and related agonists as compared to mouse L5 ET neurons. This could be of great clinical significance and underscores the importance of direct investigation of human neuron physiology and neuromodulation.

In addition to the cross-species differences that we could observe, there are undoubtedly other differences that were not accessible to the methods used in this study. Chief among these is the somato-dendritic distribution of ion channels. Previous reports have shown that the human L5 dendrites are more electrically compartmentalized compared with rat dendrites (Beaulieu-Laroche et al., 2018). This enhanced compartmentalization appears to be due in part to cross-species differences in the density of ion channels. Additionally, in the supragranular layers, human pyramidal neurons possess dendritic non-linearities not previously observed in rodent neurons (Gidon et al., 2020), that may be due to differences in the somato-dendritic expression of select ion channels. Furthermore, there could be differences in the passive membrane properties of human L5 ET neurons, or in related dendritic morphological properties (Deitcher et al., 2017; Eyal et al., 2016).

Finally, our general strategy of investigating L5 ET neurons could serve as a roadmap for studying human and non-human primate L5 ET neuron types across cortical areas. The transcriptome of L5 ET neurons varies greatly across cortical areas in both rodents and humans (Hodge et al., 2020; Tasic et al., 2018), suggesting that areal signatures in functional properties may also greatly vary as a direct consequence. Most strikingly, a few famous morphological types of L5 ET neurons are found in primate but not in rodent neocortex, including several gigantocellular neurons (e.g. the Betz cell of primary motor cortex, the Meynert cell of V1 and the Von Economo neuron of frontoinsular/anterior cingulate cortex; Allman et al., 2010; Jacobs et al., 2017). We recently utilized a similar approach to highlight conserved and divergent features of primate Betz cells and mouse corticospinal neurons (Bakken et al., 2020). Determining how cross-areal and cross-species variability in gene expression translates to phenotypic diversity at the level of cell types promises to deepen our understanding of conserved and divergent aspects of neocortical brain function through evolution and to improve prospects for translational neuroscience.

## Acknowledgments

We wish to thank the Allen Institute founder, Paul G. Allen, for his vision, encouragement and support. We also wish to thank Luke Esposito, Julie Nyhus and the Tissue Procurement team, Nick Dee, Tamara Casper, Eliza Barkan, Matthew Kroll, Herman Tung, Josef Sulc and Kirsten Crichton of the Tissue Processing Team, and the Facilities team for help in coordinating the logistics of human surgical tissue collection, transport and processing. We are also grateful to our collaborators at the local hospital sites, including Caryl Tongco, Jae-Guen Yoon, Nathan Hansen (Swedish Medical Center), Gina DeNoble and Allison Beller (Harborview Medical Center), Erica Melief, Lisa Keene, Desiree Marshall, and Caitlin Latimer (UW Medical Center) for assistance with various logistics including patient consent, case planning and coordination, and surgical tissue collection from the operating rooms. We thank Ximena Opitz-Araya, Miranda Walker and Tae Kyung Kim for molecular cloning and packaging of AAV vectors, as well as Ali Cetin, Shenqin Yao, Marty Mortrud, and Thomas Zhou of the Viral Technology team for AAV packaging. We thank Peter Chong for reagent prep and assistance with tissue processing. We thank Krissy Brouner, Augustin Ruiz, Tom Egdorf, Amanda Gary, Michelle Maxwell, Alice Pom and Jasmine Bomben of the Histology team for biocytin staining. We thank Nadezhda Dotson, Rachel Enstrom, Madie Hupp, Lydia Potekhina, Kiet Ngo, Samuel Dingman Lee, Melissa Gorham, Fiona Griffin, Eric Lee, and Shea Ransford of the Imaging team for imaging of biocytin filled cells. We thank Darren Bertagnolli, Michael Tieu, Delissa McMillen, Thanh Pham, Christine Rimorin, Katelyn Ward, Alexandra Glandon, and Amy Torkelson of the RSeq core for scRNA-seq processing. We thank Jeremy A. Miller, Osnat Penn, and Zizhen Yao for contributions to the Patch-seq tree-based mapping algorithms, and Jeff Goldy and Olivia Fong for Patch-seq data management and updates in MolGen Shiny. We thank Brian Lee, Jim Berg, Lindsay Ng, Rusty Mann, Jessica Trinh and other members of the Ephys Core for assistance with sample processing and reporting. We thank the Animal Care Team for mouse husbandry, the Allen Institute Transgenic Colony Management team for colony management, the Allen Institute Laboratory Animal Services team for preparation and delivery of experimental mice. We thank Abi Gibson for help with morphological reconstructions.

## Funding

This work was funded by the Allen Institute for Brain Science and also supported in part by the U.S. National Institutes of Health (NIH) grant U01 MH114812-02 to E.S.L., NIH BRAIN Initiative award RF1MH114126 from the National Institute of Mental Health to E.S.L., J.T.T., and B.P.L., NIH BRAIN Initiative award RF1MH121274 to B.T. and H.Z, NIH grants P51OD010425 from the Office of Research Infrastructure Programs (ORIP), NIA grant AG005136 to the UW ADRC Neuropathology Core, a grant from the Nancy and Buster Alvord Endowment to C.D.K. and UL1TR000423 from the National Center for Advancing Translational Sciences (NCATS). Its contents are solely the responsibility of the authors and do not necessarily represent the official view of NIH, ORIP, NCATS, the Institute of Translational Health Sciences or the University of Washington National Primate Research Center.

## Author contributions

Conceptualization and management of the project: B.E.K., E.S.L. and J.T.T.; Patch-seq and dendritic recording and analysis: B.E.K.; mFISH data generation and analysis: R.D.H., A.M.Y.; RNA-seq data generation and analysis: K.S., R.D.H, N.L.J., T.E.B., S.O., B.T.; histology: M.M., R.DF.; imaging: P.R.N., R.DF.; neuron 3D reconstruction and analysis: R.DF., S.A.S., R.D., M.M.; data visualization tools: L.T.G, B.T.; AAV vectors: T.L.D., J.T.T.; neurosurgery and human surgical tissue acquisition: R.P.G., D.L.S., C.C., J.G.O., A.L.K., C.D.K.; manuscript preparation: B.E.K., S.O, and J.T.T. with input from all authors. Program leadership: E.S.L., H.Z., and C.K.

## EXPERIMENTAL MODEL AND SUBJECT DETAILS

### Human surgical specimens

Surgical specimens were obtained from local hospitals (University of Washington Medical Center, Swedish Medical Center and Harborview Medical Center). Hospital institute review boards approved all procedures involving human tissue before commencing the study and all patients provided informed consent. Data included in this study were obtained from neurosurgical tissue resections for the treatment of refractory temporal lobe epilepsy (n=23) or deep brain tumors (n=3) in 21 male and 5 female patients with a mean age of 39.23 ± 3.07 years (Table S1).

### Mouse specimens

All procedures involving mice were approved by the Allen Institute’s Institutional Care and Use Committee. Mixed strains of male and female transgenic mice, from 60-90 days old were used for experiments. Mice were maintained on a 12-hour light/dark cycle in a temperature and humidity controlled room. Mice were housed 3-5 per cage with free access to food and water.

### Macaque specimens

All procedures involving macaque monkeys were approved by the University of Washington’s Institutional Care and Use Committee. Male (n=2) and female (n=1) macaques (macaca nemestrina) from 3-17 years old designated for use in the Washington National Primate Research Center’s Tissue Distribution Program were used for experiments. Monkeys were housed in individual cages on a 12-hour light/dark cycle in a temperature and humidity controlled room.

## METHOD DETAILS

### Acute brain slice preparation

Brain slice preparation was similar for all species. Human neurosurgical specimens deemed not to be of diagnostic value were placed in a sterile, prechilled, carbogenated (95% O_2_/5% CO_2_) container filled with an artificial cerebrospinal fluid composed of (in mM): 92 N-methyl-D-glucamine (NMDG), 2.5 KCl, 1.25 NaH_2_PO_4_, 30 NaHCO_3_, 20 4-(2-hydroxyethyl)-1-piperazineethanesulfonic acid (HEPES), 25 glucose, 2 thiourea, 5 Na-ascorbate, 3 Na-pyruvate, 0.5 CaCl_2_**□**4H_2_O and 10 MgSO_4_**□**7H_2_O. Surgical specimens were quickly transported from the surgical site to the Institute while continuously bubbled with carbogen. Macaques were anesthetized with sevoflurane gas during which the entire cerebrum was extracted and placed in the same aCSF described above.

After brain extraction, monkeys were administered intravenous injection of a lethal dose of sodium-pentobarbital. We then dissected the superior temporal gyrus of the temporal lobe for brain slice preparation. Mice were deeply anesthetized by intraperitoneal administration of Avertin (20 mg/kg) and were perfused through the heart with the same NMDG aCSF described above.

Brain specimens were sectioned on a Compresstome VF-200 using a zirconium ceramic blade (Precisionary Instruments) at 300 or 350 μm using the protective recovery method (Ting et al., 2014). To ensure that the dendrites of pyramidal neurons were relatively intact, macaque and human specimens were trimmed and mounted such that the angle of slicing was perpendicular to the pial surface. Mouse brains were sectioned in the coronal plane.

### Patch clamp recordings

Brain slices were placed in a submerged, heated recording chamber that was continuously perfused with carbogenated aCSF consisting of (in mM): 119 NaCl, 2.5 KCl, 1.25 NaH2PO4, 24 NaHCO3, 12.5 glucose, 2 CaCl2·4H2O and 2 MgSO4·7H2O (pH 7.3-7.4). Slices were visualized with an Olympus BX51WI microscope and infrared differential interference contrast (IR-DIC) optics and a 40x water immersion objective.

Patch pipettes (2-5 MΩ for somatic; 4-8 MΩ for dendritic) were filled with one of two internal solutions. The first solution contained (in mM): 126.0 K-gluconate, 10.0 HEPES, 0.3 EGTA, 4.0 mM KCl, 4 Mg-ATP, 0.3 Na2-GTP, 10.0 phosphocreatine disodium salt hydrate, 0.5% biocytin and .02 Alexa 594 or 488. The second pipette solution was used for Patch-seq experiments and contained (in mM): 110.0 K-gluconate, 10.0 HEPES, 0.2 EGTA, 4 KCl, 0.3 Na2-GTP, 10 phosphocreatine disodium salt hydrate, 1 Mg-ATP, 20 μg/ml glycogen, 0.5U/μL RNAse inhibitor (Takara, 2313A), 0.5% biocytin and 0.02 Alexa 594 or 488. The pH of both solutions was adjusted to 7.3 with KOH. Alexa filled neurons were visualized upon termination of recording using a 540/605 nm excitation/emission filter set. The liquid junction potential was calculated to be −13 mV and was not corrected.

Whole cell somatic and dendritic recordings were acquired using a Multiclamp 700B amplifier and either PClamp 10 data acquisition software or custom MIES acquisition software (https://github.com/AllenInstitute/MIES/) written in Igor Pro. Electrical signals were digitized at 20-50 kHz by a Axon Digidata 1550B (Molecular Devices) or a ITC-18 (HEKA) and were filtered at 2-10 kHz. Upon attaining whole cell current clamp mode, the pipette capacitance was compensated and the bridge was balanced. Access resistance was monitored throughout the recording and was 8-30 MΩ for somatic recordings and 15-40 MΩ for dendritic recordings.

### Processing of Patch-seq samples

Prior to data collection for these experiments, all surfaces were thoroughly cleaned with RNAse Zap, and as needed DNAse Away. At the end of Patch-seq recordings, negative pressure (~−20 mbar) was applied through the pipette for ~5 minutes after which the nucleus was extracted by very slow pipette withdrawal with higher negative pressure (~−70 to −100 mbar). The pipette was removed from the recording chamber and the contents of the pipette were expelled into a PCR tube containing lysis buffer (Takara, 634894). Patch-seq sample tubes were held on dry ice in a benchtop plexiglass enclosure throughout the recording session to ensure collected samples remained free of RNAse and DNAse contamination. Sample tubes were then transferred to −80C for storage until further processing. cDNA libraries were produced using the SMART-Seq v4 Ultra Low Input RNA Kit for Sequencing (Takara 634894, Lot 1709695A) according to the manufacturer’s instructions, using 20 PCR cycles for cDNA amplification. Samples proceeded through library construction using Nextera XT DNA Library Preparation Kit (Illumina FC-131-1096) according to the manufacturer’s instructions except at 0.2x reaction size. Samples were sequenced to approximately 1million paired-end 50b reads//sample.

### Biocytin histology

As described previously (Gouwens et al., 2019), a horseradish peroxidase (HRP) enzyme reaction using diaminobenzidine (DAB) as the chromogen was used to visualize biocytin-filled neurons following physiological recordings. 4,6-diamidino-2-phenylindole (DAPI) stain was also used to identify cortical layers.

Slices were mounted and imaged on an upright AxioImager Z2 microscope (Zeiss, Germany) equipped with an Axiocam 506 monochrome camera and 0.63x optivar. High-resolution image stacks were captured with a 63X objective at 0.44 μm increments along the Z axis. ZEN software was used to stitch tiled images.

### Multiplex fluorescence in situ hybridization

Fresh-frozen human (postmortem and surgical) MTG, macaque (*M. nemestrina)* MTG, and mouse TEa tissues were sectioned at 14-16 μm onto Superfrost Plus glass slides (Fisher Scientific). Sections were prepared from at least two donors per species. Sections were dried for 20 minutes at −20°C and then vacuum sealed and stored at −80°C until use. The RNAscope multiplex fluorescent v1 kit was used per the manufacturer’s instructions for fresh-frozen tissue sections (ACD Bio), except that fixation was performed for 60 minutes in 4% paraformaldehyde in 1X PBS at 4°C, post-dehydration drying was done for 15 minutes at 37°C and protease treatment was shortened to 10 minutes. Sections were imaged using a 60X oil immersion lens on a Nikon TiE fluorescence microscope equipped with NIS-Elements Advanced Research imaging software (version 4.20). For all RNAscope mFISH experiments, positive cells were called by manually counting RNA spots for each gene. Cells were called positive for a gene if they contained ≥ 3 RNA spots for that gene. Lipofuscin autofluorescence was distinguished from RNA spot signal based on the larger size of lipofuscin granules and broad fluorescence spectrum of lipofuscin.

To quantify the fraction of putative ET cells (defined as *FAM84B* and *SLC17A7* double positive cells) in layer 5 in each species, the boundary of layer 5 was first delineated using DAPI to identify cortical layers. The total number of *SLC17A7*+ cells within layer 5 was quantified and then the total number of *SLC17A7*+,*FAM84B*+ cells in layer 5 was quantified. The percentage of putative ET cells (*SLC17A7*+, *FAM84B+*) was then calculated as a fraction of the total number of *SLC17A7*+ cells in layer 5. Counts were repeated on at least 2 donors per species and at least 2 sections per donor.

## QUANTIFICATION AND STATISTICAL ANALYSIS

### Neurophysiology

We used three basic current stimuli to probe the intrinsic membrane properties of L5 neurons. In the first protocol, we measured the voltage response to a series of 1s steps from −150 pA to +50 pA in +20 pA increments. Maximum and steady-state input resistance (R_N_) were calculated from the linear portion of the current-maximum or steady state voltage relationship generated in response to these current injections. Voltage sag was defined as the ratio of maximum to steady-state R_N_. Rebound slope was calculated from the slope of the rebound amplitude as a function of steady-state membrane potential.

The second stimulus was a chirp stimulus that increased in frequency either linearly from 1-15 Hz over 15 s or logarithmically from 0.2-40 Hz over 20s. The amplitude of the chirp was adjusted for each neuron to produce a peak-to-peak voltage deflection of ~10 mV. Impedance amplitude (ZAP) was derived as the ratio of the Fourier transform of the voltage response to the Fourier transform of the chirp: 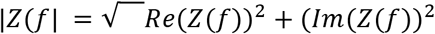,where Im(Z(f)) and Re(Z(f)) are the imaginary and real parts of the impedance Z(f), respectively. The frequency at which the maximum impedance occurred was the resonant frequency (*f*_R_). Resonance strength (Q) was measured as the ratio of the maximum impedance amplitude to the impedance amplitude at 1 Hz. The 3dB cutoff was defined as the frequency at which the ZAP profile attenuated to a value of 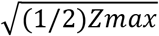. Impedance phase (ZPP) was derived as: 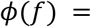 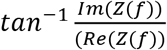. Total inductive phase (Zϕ) was defined as the area under the inductive part of the ZPP. Synchrony phase was the frequency at which the ZPP was 0.

The third stimulus was a series of 1 second depolarizing current injections increasing in amplitude by +50 pA/step. We measured the number of spikes and the first instantaneous firing rate in response to each current injection. Gain was defined as the linear slope of this action potential frequency-current relationship. Single action potential properties were measured from voltage responses to the lowest current that produced a spike. Action potential threshold was defined as the voltage at which the first derivative of the voltage response exceeded 20 V/s. AP width was measured at half the amplitude between threshold and the peak voltage. Fast AHP was defined relative to threshold. Spike frequency accommodation (SFA) and medium afterhyperpolarization (mAHP) were calculated from current injections producing ~10 spikes during the 1 s step. SFA was defined as the ratio of the second to the last interspike interval. The mAHP was defined as the minimum voltage after the spike train. For dendritic recordings we also quantified the area and width of plateau potentials. The threshold for a plateau was defined as the voltage where the first derivative reached 2 V/s. Plateau width at half maximum amplitude and plateau area were calculated relative to this threshold.

We grouped neurons based on physiological properties derived from these current injections using hierarchical clustering. First we performed principal component analysis on all physiological features. Principal components explaining at least 1% of the variance were then used to cluster neurons into groups using Ward’s algorithm. The number of clusters was determined using the sigClust package in R which generates a p value for the null hypothesis that data points are drawn from a single Gaussian as opposed to two Gaussian distributions. The p value for this analysis was set at < 0.01, which yielded three clusters.

Random forest classifiers were constructed using the randomForest package in R (Liaw and Wiener, 2002). We varied the percentage of cells included in the training data set from 10-50%. For each training set, we constructed 100 forests consisting of 100 trees. Mean accuracy and importance values were calculated from each set of forests on the data held out of the training sets.

Comparisons across groups are presented as geometric box plots unless otherwise denoted. Statistical significance was assayed using ANOVAs, t-tests, Mann-Whitney U tests, or Person’s correlation where appropriate. P values were FDR adjusted for multiple comparisons. Effect sizes are reported as eta-squared values.

### Morphological reconstruction

Dendritic reconstructions were generated based on a digital 3D image stack that was run through a Vaa3D-based image processing and reconstruction pipeline, as previously described (Gouwens et al., 2019). Somatic morphology was quantified directly via a maximum projection of a series of 63x images of biocytin and DAB reacted neurons. The initial apical shaft width was measured 50 μm from the center of the soma.

### Patch-Seq sample mapping

To determine the corresponding transcriptomic cell type for each Patch-seq sample, we utilized a tree-based mapping strategy. For each neuron, we computed its correlation with each branch of the reference cell type taxonomy, starting from the root and working towards the leaves. Marker genes associated with each branch of the taxonomy were used for correlations. The confidence of the mapping was determined by applying 100 bootstrapping iterations of the process. For each iteration 70% of the reference cells and 70% of maker genes were randomly sampled for mapping. The percentage of times a given cell mapped to a given transcriptomic cell types was the mapping probability. Only neurons with a mapping probability greater than 50% to a given terminal leaf were included. As an additional quality control measure, only Patch-seq samples with a normalized summed expression of “on”-type marker genes (NMS; (Tripathy et al., 2018) were included. We mapped mouse Patch-seq samples to a published mouse VISp scRNA-seq cell type taxonomy (Tasic et al., 2018) and human samples to a published human MTG snRNA-seq cell type taxonomy (Hodge et al., 2019).

### Differential Gene Expression analysis

To identify differentially expressed genes between L5 ET and L5 IT neurons for each species, expression matrices were trimmed to only include genes with one-to-one orthologs in human and mouse (downloaded from NCBI Homologene in November, 2019). Raw expression matrices for each brain region were CPM normalized then Log2 transformed. Average expression was determined for each subclass (i.e. L5 ET and L5 IT), then log fold change was determined between subclasses for each brain region. The following clusters from human MTG were used for L5 IT subclass input; Exc L5-6 RORB TTC12, Exc L4-5 RORB FOLH1B, Exc L5-6 THEMIS C1QL3, Exc L4-6 RORB SEMA3E, and Exc L4-6 RORB C1R; and the Exc L4-5 FEZF2 SCN4B cluster was used as input for L5 ET. For mouse, all previously defined L5 IT and L5 PT clusters were used as input. Initial GO analysis of conserved L5 ET DE genes (> 0.5 Log2FC across all three brain regions) with PANTHER Classification System (Mi et al., 2019) revealed numerous significant GO biological process categories for axonogenesis, axon guidance, axon development, etc. Additionally, numerous synaptic-related categories were enriched (i.e synaptic membrane adhesion, synaptic organization, synaptic structure, etc.). Respective terms were aggregated into an Axon Guidance gene list, and a Synaptic Regulation gene list for visualization purposes in Figure 1, with aggregated gene lists shown in Table S2. Human L5 ET-enriched (> 1 Log2FC) DE genes from each GO category were plotted as line plots and colored red if L5 ET neurons had > 1 Log2FC expression over L5 IT in all brain regions, indicating conserved L5 ET DE genes. Lines were colored blue if L5 ET expression was > 1 Log2FC in human MTG and < 0 in both mouse regions, indicating human-specific L5 ET DE genes.

---

**Figure S1.**
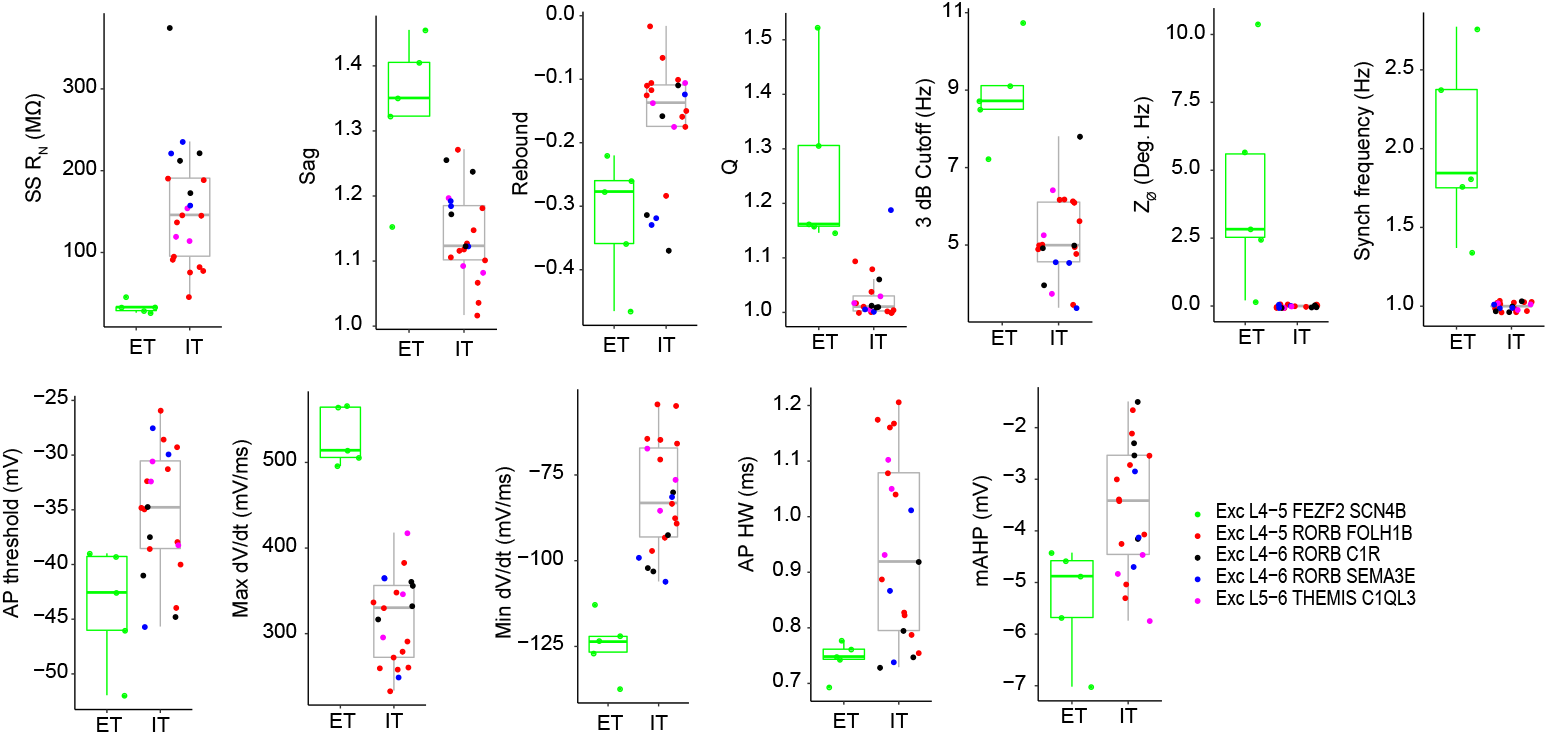
Additional pairwise differences in the intrinsic membrane properties of transcriptomically defined cell types. All comparisons were * p < 0.05, FDR corrected Mann-Whitney U test.

**Figure S2.**
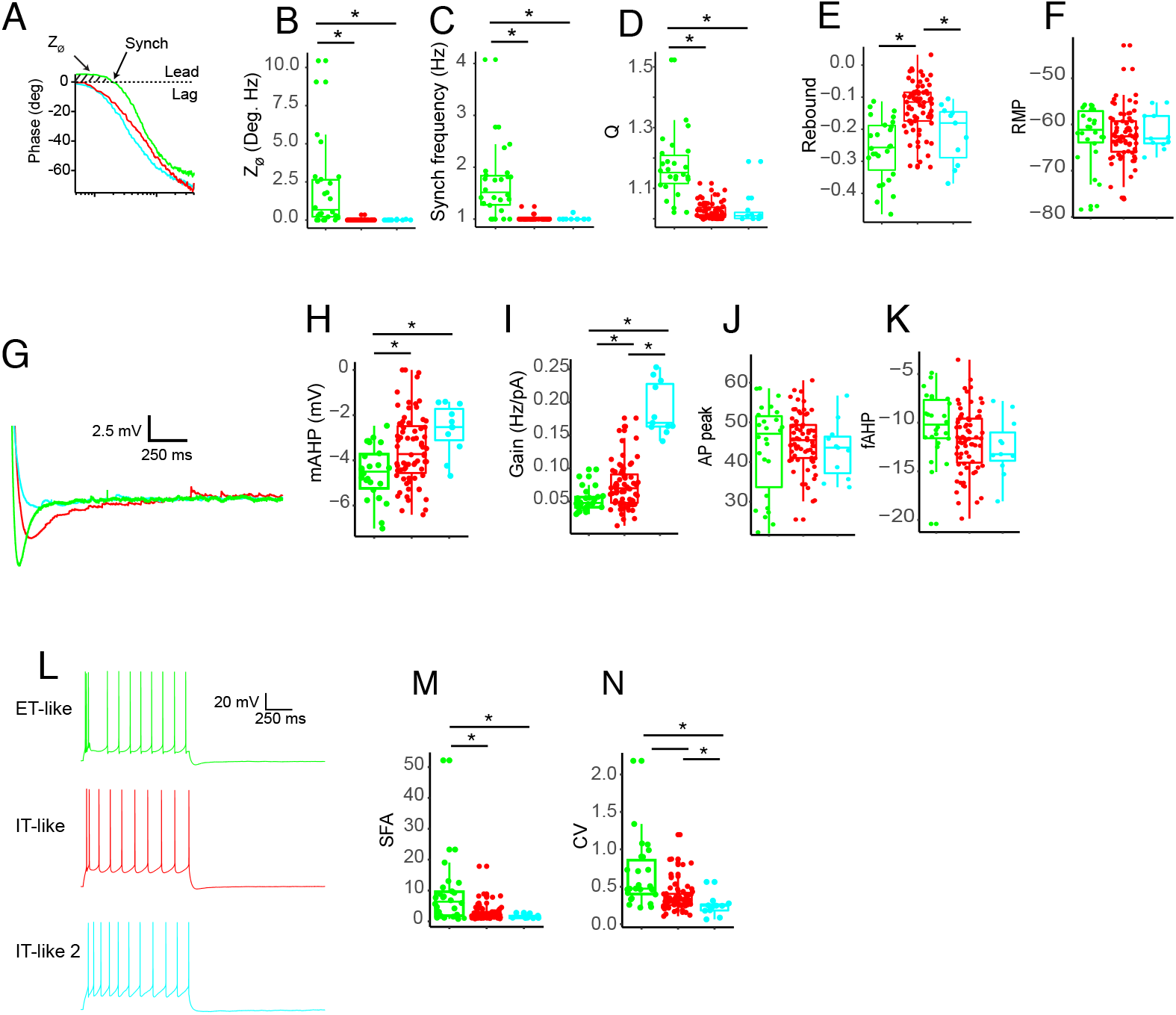
Additionai pairwise comparisons of intrinsic membrane properties of physiologically defined L5 pyramidal neurons in human middle temporal gyrus. A) Example impedance phase plot for ET-like, IT-like 1 and IT-like 2 neurons. Dashed line indicates the frequency at which the current and voltage are in phase with one another (ZPP). B-F) Additional pairwise comparisons of subthreshold properties. G) Example medium afterhyperpolarization produced by ~10 Hz firing. H-K) Additional pairwise comparisons of suprathreshold properties. L) Example ~10 Hz firing. Pairwise comparison of M) spike frequency accommodation and N) Coefficient of variation. * p < 0.05, FDR corrected Mann-Whitney U test.

**Figure S3.**
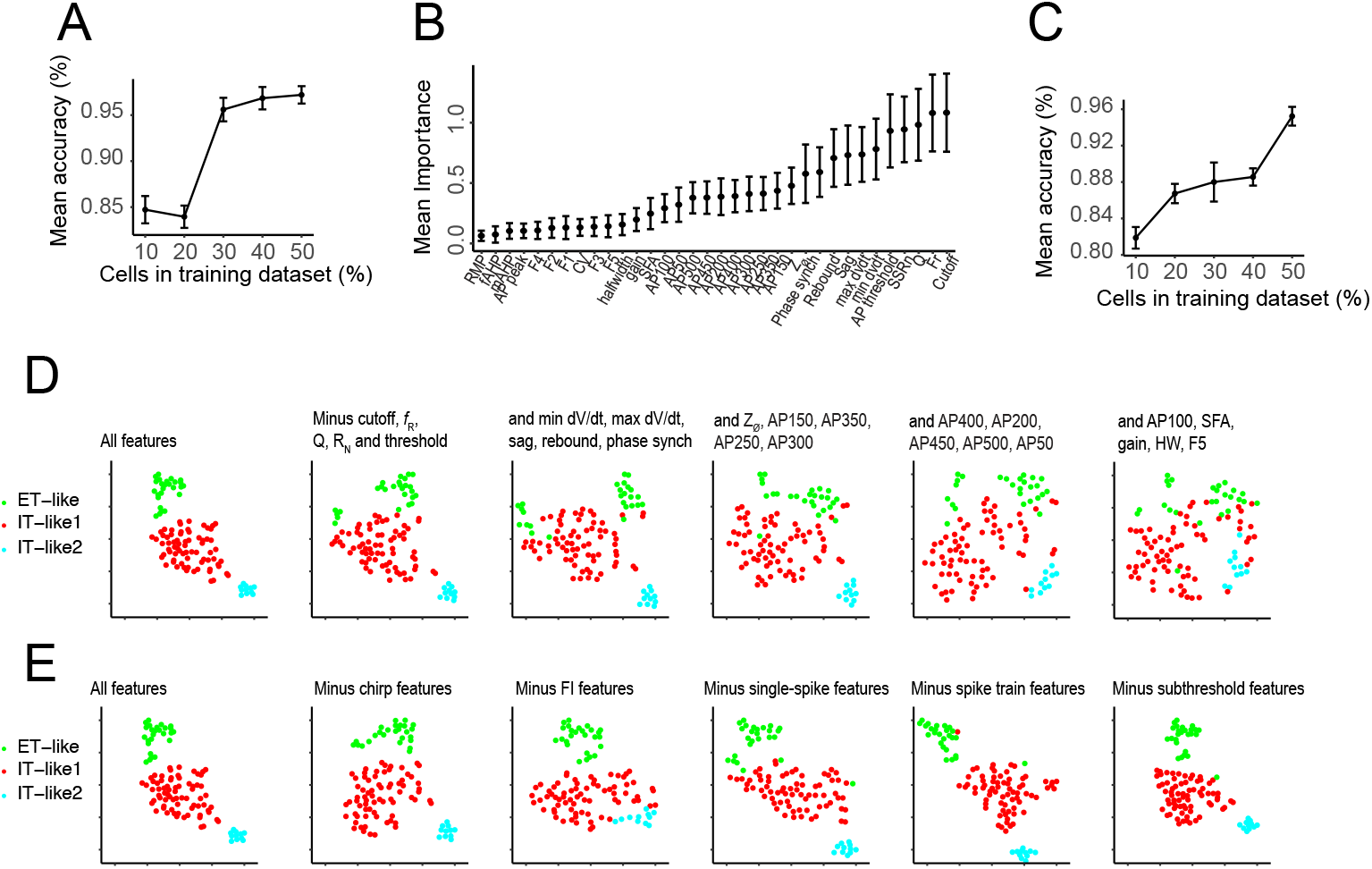
The distinctiveness of L5 neuron properties is determined by an aggregate of physiological features. A) Performance of a random forest classifier as a function of the percentage of cells included in the training dataset. B) Mean importance of physiological features as determined by random forest classifier. C) Performance of random forest classifier using only the top 10 important features. D)tSNE projections based on all physiological features (left) and after subsets of features are successively removed. Features were removed in order based on their mean importance determined by the random forest classifier. E) tSNE projections based on all features (left) and where specific features are removed as indicated.

**Figure S4.**
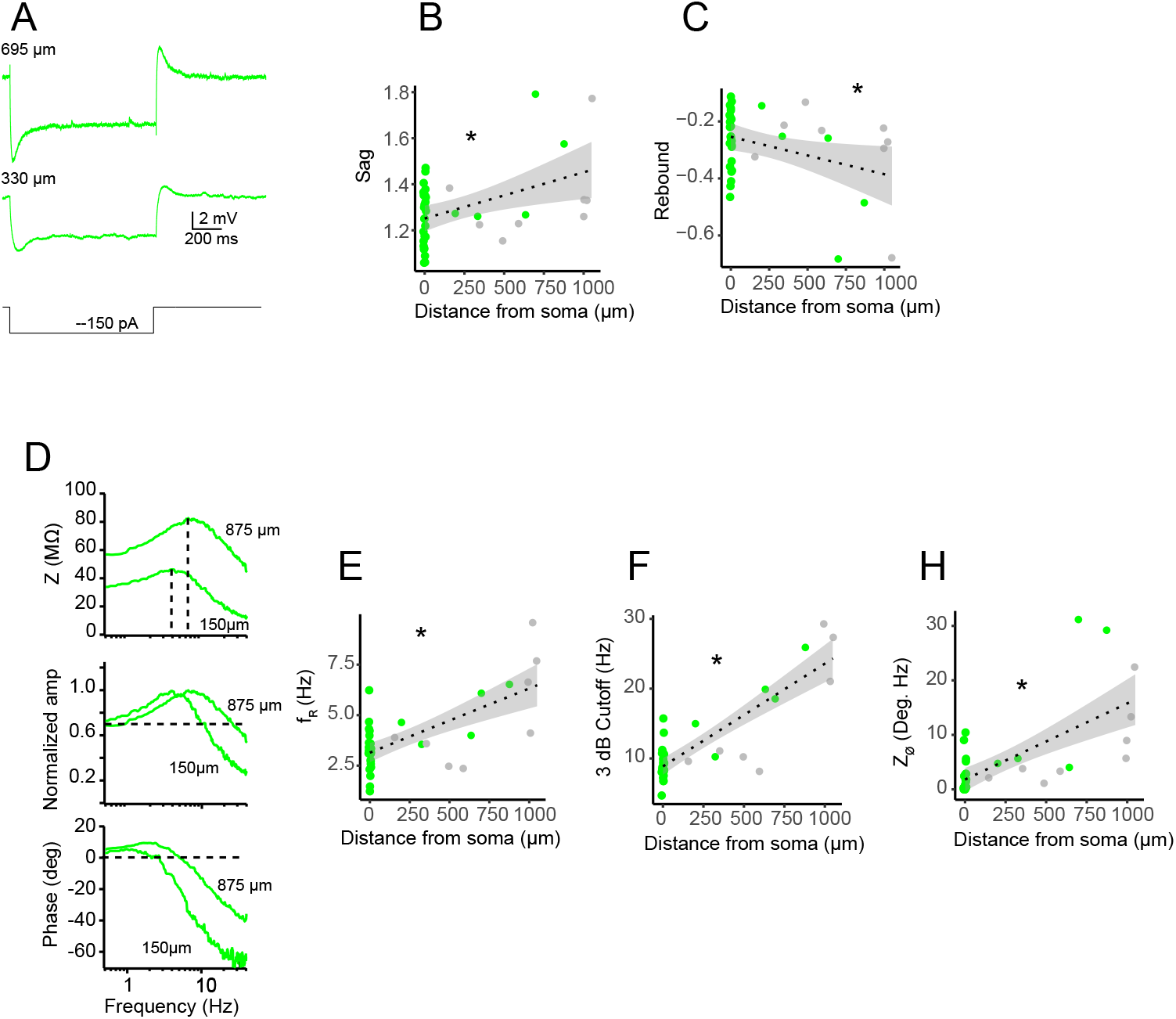
Intrinsic membrane properties of putative L5 ET neuron dendrites. A) Voltage response to hyperpolarizing current injection recorded at two different dendritic recording sites. B) Sag (p = 0.008) and C) rebound potentials (p = 0.04) as a function of distance from soma. D) Example ZAP (top), normalized frequency response (middle) and ZPP (bottom) for dendritic recordings. E) Resonance frequency, F) 3dB cutoff (p < 0.001) and H) total inductive phase (p < 0.001) as a function of distance from soma, p values are for FDR corrected Pearson’s correlation.

**Figure S5.**
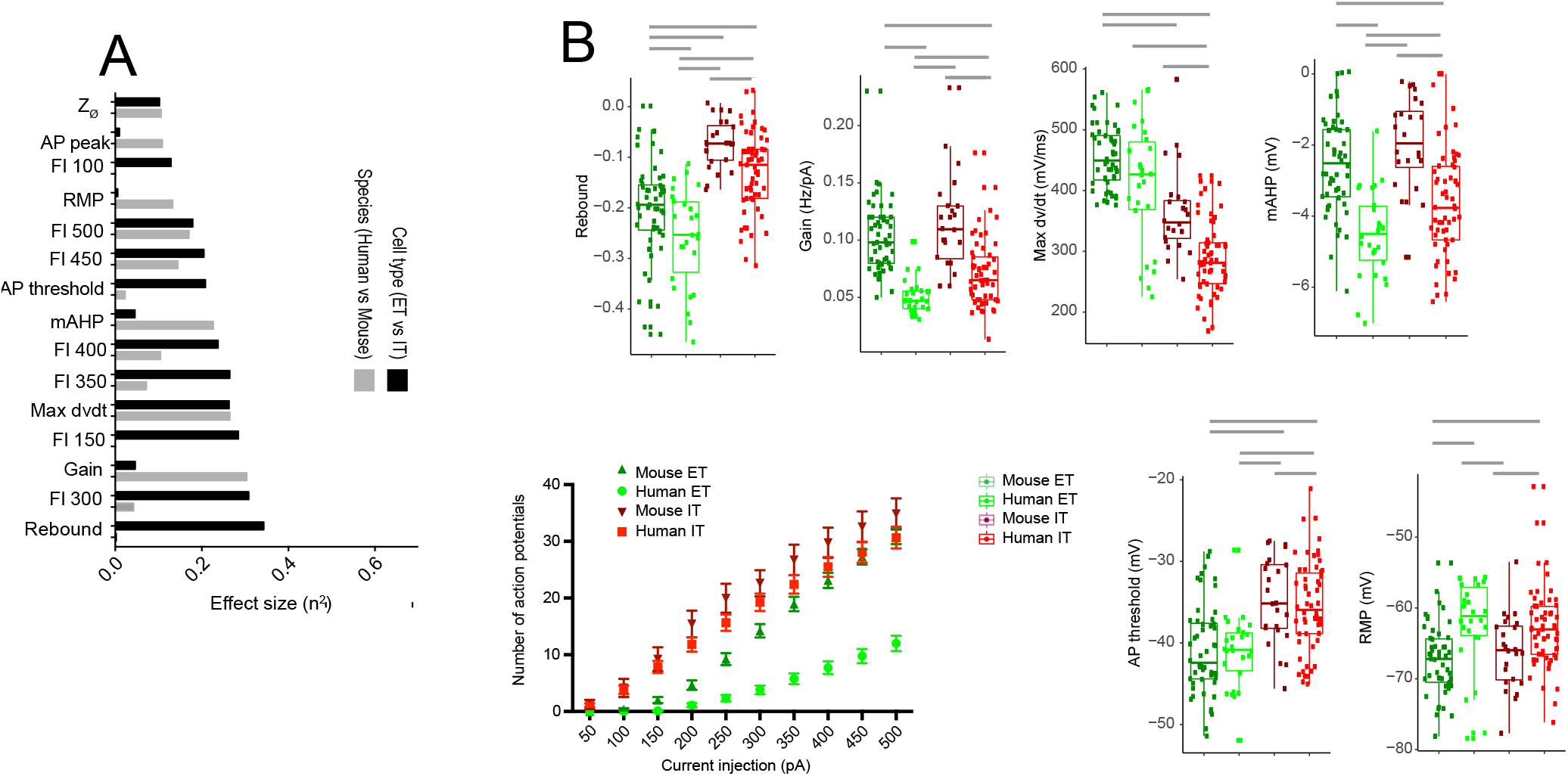
Additionai cross-species/cell type comparisons. A) The next 15 largest effect sizes resulting from ANOVA with cell type (ET versus IT) and species (human versus mouse) as the factors. B) Physiological features plotted as a function of species and cell type. Gray bars denote * p < 0.05, FDR corrected Mann-Whitney U test.

## Notes

### Competing Interest Statement

The authors have declared no competing interest.

